# Repurposing mitochondrial-targeting anthelmintic agents with GLUT1 inhibitor BAY-876 for cancer therapy

**DOI:** 10.1101/2024.05.02.592272

**Authors:** Tanner J. Schumacher, Ananth V. Iyer, Jon Rumbley, Conor T. Ronayne, Venkatram R. Mereddy

## Abstract

Cancer cells alter their metabolic phenotypes with nutritional change. Single agent approaches targeting mitochondrial metabolism in cancer have failed due to either dose limiting off target toxicities, or lack of efficacy *in vivo*. To mitigate these clinical challenges, we investigated the potential utility of repurposing FDA approved mitochondrial targeting anthelmintic agents, niclosamide and pyrvinium pamoate, to be combined with GLUT1 inhibitor BAY-876 to enhance the inhibitory capacity of the major metabolic phenotypes exhibited by tumors. To test this, we used breast cancer cell lines MDA-MB-231 and 4T1 which exhibit differing basal metabolic rates of glycolysis and mitochondrial respiration, respectively. Here, we found that specific responses to mitochondrial and glycolysis targeting agents elicit responses that correlate with tested cell lines basal metabolic rates and fuel preference, highlighting the potential to cater metabolism targeting treatment regimens based on specific tumor nutrient handling. Inhibition of GLUT1 with BAY-876 potently inhibited glycolysis in both MDA-MB-231 and 4T1 cells, and niclosamide and pyrvinium pamoate perturbed mitochondrial respiration that resulted in potent compensatory glycolysis in the cell lines tested. In this regard, combination of BAY-876 with both mitochondrial targeting agents resulted in inhibition of compensatory glycolysis and subsequent metabolic crisis. These studies warrant further investigation into targeting tumor metabolism as a combination treatment regimen that can be tailored by basal and compensatory metabolic phenotypes.

## INTRODUCTION

Cells undergo catabolic and anabolic metabolism based on their specific needs whether it be to accumulate energy or initiate replication. These processes occur typically in a homeostatic manner, regulated through signaling or via feedback loops. A quiescent cell uptakes carbon-based nutrients from their environment fated for oxidative catabolism in which electrons will get transferred to NAD^+^ or FAD, generating NADH and FADH_2_, respectively. The electrons will pass through the electron transport chain (ETC) to reduce O_2_ to water to generate ATP for energy. A replicating cell will not only undergo the acquisition of energy from carbon catabolism, but due to various pro-proliferation signaling, will redirect nutrients towards anabolic reactions to generate biomass required to produce a daughter cell. This continuously active anabolic state will eventually pull all the available local nutrients required for this anabolic state. Normal cells will sense the depletion of local nutrients and turn off proliferative signaling through a variety of metabolic feedback loops.^1–4^ In response to nutrient-rich conditions, anabolic processes including glycolysis, lipid, nucleotide, and protein synthesis will be activated with concomitant downregulation of autophagy.^1–7^ Conversely, low nutrient availability will decrease anabolic signaling and increase catabolic signaling pathways.^1–7^ Cancer cells undergo metabolic reprogramming that allow them to maintain a proliferative state and bypass metabolic feedback loops or hyperactivate proliferative signaling. Such metabolic signaling has rendered targeting metabolic pathways as single agents clinically ineffective as activation of compensatory metabolism circumvents antitumor efficacy in these trials.^8^

The metabolism of tumors is in constant equilibrium between glycolysis and oxidative phosphorylation (OxPhos) and is dictated by gradients of nutrients and oxygen. Interestingly, many utilize glycolytic metabolism even in the presence of sufficient oxygen (Warburg effect, aerobic glycolysis), an oncogenic process that has been leveraged as a therapeutic target.^9–12^ Recent literature has demonstrated that mitochondria remain functional even in cases where aerobic glycolysis is occurring as a means to offer biosynthetic intermediates to anabolic-proliferating cells while serving as sources of bioenergetics to cells in oxygenated environments.^13–15^ Thus, targeting both glycolysis and mitochondrial respiration may provide a more durable treatment strategy when compared to targeting a single pathway.

The maintenance of the proton gradient across the inner mitochondrial membrane allows for a biophysical means for pharmacological targeting. One class of mitochondrial targeting agents, protonophores, are chemical entities that can carry a proton across the inner mitochondrial membrane, deliver a proton, and then cycle back across the membrane to the inner mitochondrial space. Bringing protons into the mitochondrial matrix serves to uncoupling ATP synthesis from the proton gradient, destabilizing or inhibiting OxPhos. Delocalized lipophilic cations (DLCs) are positively charged molecules that can distribute that charge over a large hydrophobic area to allow them to cross membranes.^16,17^ Due to the negative membrane potential (ΔΨ_m_) across the inner mitochondrial membrane, DLCs tend to accumulate in the mitochondria matrix.^16,17^ Due to the importance of mitochondrial respiration in cancer, protonophores, DLCs and other mitochondrial targeting agents have been used as experimental tools and as therapeutic agents for a variety of diseases, including cancer. However, metabolic reprogramming observed in cancer cells provides a therapeutic opportunity to combine these mitochondrial targeting agents with a glycolysis inhibiting agent.

There are currently several thousand FDA approved and clinically available drugs for a wide variety of ailments and diseases having undergone extensive toxicity and pharmacokinetic/pharmacodynamic analysis in humans with often well-reported side effects and contraindications. Although these drugs are in the clinical setting for a particular disease, their biological target and/or mechanism can be found to be relevant in a novel disease context. Hence, many older drugs can be qualified as suitable candidates for repurposing for new indications, principally for cancer. In the past decade, increasing interest in leveraging niclosamide as an anticancer agent has led to the discovery that niclosamide targets a wide range of pathways such as regulation of Wnt/β-catenin^18,19^, mTORC1^20–22^, STAT3^23–25^, NF-κB^25,26^, and Notch signaling pathways.^27,28^ Several literature reports indicate niclosamide’s activity in several cancers such as adrenocortical carcinoma^29^, head and neck cancer^30^, colon cancer^31^, leukemia^32^, lung cancer^33^, glioblastomas^34^, renal cell carcinoma^35^, prostate cancer^36^, ovarian cancer^37^, and breast cancer^38^. Pyrvinium pamoate has also been utilized as an anticancer agent due to its ability to modulate mitochondrial activity. Pyrvinium pamoate has been found to significantly reduce colorectal tumor spheroid growth by 62%.^19^ In a pancreatic carcinoma cell line MIAPaCa-2, pyrvinium pamoate was found to co-localize in the mitochondrial with mitotracker green where it was found to elicit a time dependent decrease in ATP levels and cell viability under glucose starvation conditions.^39^ Some work with pyrvinium pamoate was also done with ρ^ο^ cancer cell lines lacking mitochondrial DNA where it was found to accumulate in mitochondria less and displayed resistance to PP treatment.^40,41^ Some studies on isolated mitochondria have shown that pyrvinium pamoate acts as an inhibitor of complex I activity, inhibits the NADH-fumarate reductase system under hypoxic low glucose conditions, and can stimulate complex II activity under normoxic glucose positive conditions.^42–44^ In the current study, we evaluated the potential utility of repurposing clinically used anthelmintic agents niclosamide and IMD-0354 (protonophores), along with pyrvinium pamoate (DLC) for their ability to target mitochondrial metabolism and characterize their effects on compensatory glycolysis in two metabolically distinct and aggressive triple-negative breast cancer cell lines. As a proof of concept, we evaluated the ability of pharmacologically targeting mitochondrial and glycolytic metabolism by combining these clinically used agents with a preclinically validated, potent, and highly specific glucose transporter 1 (GLUT1) inhibitor, BAY-876. GLUT1 imports extracellular glucose for glycolysis, where its end product, pyruvate, can also be utilized for mitochondrial respiration. Interestingly, the GLUT1 isoform has been shown to be highly overexpressed in many solid tumors, qualifying it as a potential therapeutic target.^45^ These studies demonstrate that oncogenic transformation of basal bioenergetic phenotypes, along with compensatory metabolic capacities provide opportunities to circumvent reprogrammed metabolism with mitochondrial and glycolysis inhibitors.

## RESULTS AND DISCUSSION

It is well appreciated that cancer cells exhibit differential dependencies on metabolic pathways in the context of nutrient and oxygen availability, catabolic or anabolic nutrient handling, along with concomitant selection of oncogenes that promote or suppress metabolic fuel choice.^46–49^ We sought to examine the literature for FDA approved drugs, along with pharmacological tools, that have been shown to exhibit a wide variety of metabolism targeting capacities to evaluate their effects on cancer cell proliferation. To test this, we utilized two cancer cell lines: human triple-negative breast cancer MDA-MB-231 and murine metastatic breast cancer 4T1 as model systems derived from aggressive and difficult to treat breast cancers. Interestingly, MDA-MB-231 exhibits high basal glycolytic rates when compared to the high oxidative phosphorylation rates observed in 4T1 (**Fig. 1A**). Thus, these cell lines provide a means to test the efficacy of a diverse range of metabolism inhibitors in contexts of differing fundamental metabolic programs.

**Figure 1:**
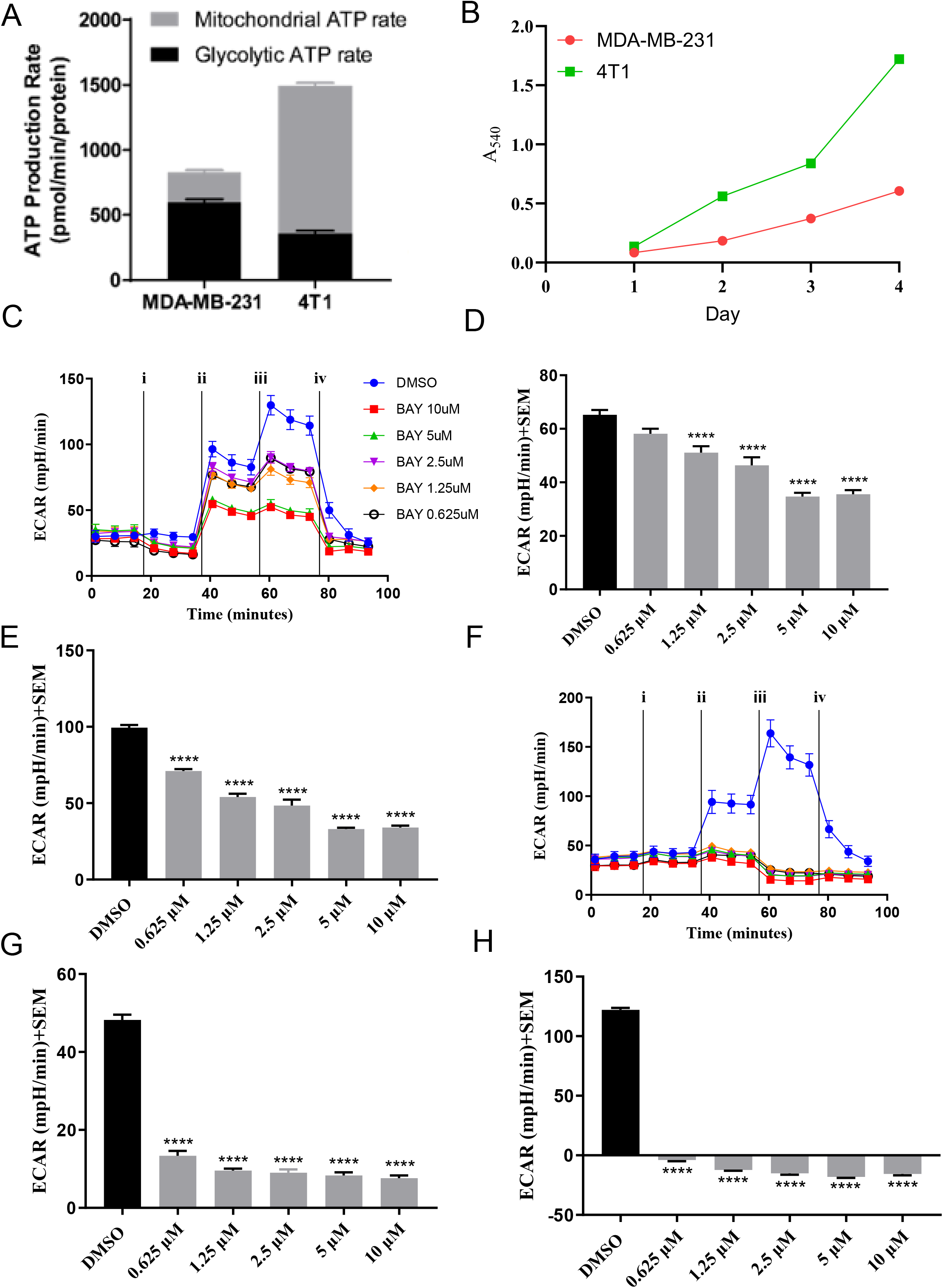
**A.** Seahorse XFe96^®^ ATP rate assay. Rates indicative of ATP production rate +SEM (pmol/min) from glycolysis (black) or mitochondria. Results are from three separate experiments and are normalized to total protein using bicinchoninic acid (BCA). **B.** Growth curve for 4T1 and MDA-MB-231 cells. Each data point represents 6 technical replicates. On each day, media was aspirated, rinsed three times with cold PBS and allowed to dry fix for a minimum of 24 hours. After all wells have dry fixed for 24 hours, 0.5% w/v SRB in 1% acetic acid was added to the wells for 45 minutes, aspirated and rinsed three times with 1% acetic acid, allowed to dry, solubilized in 10mM Tris bas (pH 10.2) and absorbance was measured at 540 nm. **C.** MDA-MB-231 extracellular acidification rates (ECAR (mpH/min)) vs. time after injection of i) treatment or DMSO ii) glucose iii) oligomycin and iv) 2-deoxyglucose. **D.** calculated ECAR after addition of glucose which indicates extent of cellular glycolysis and inhibition from treatment of BAY-876 (10-0.625µM) in MDA-MB-231 cells **E.** ECAR indicating glycolytic capacity after uncoupling mitochondrial ATP synthesis after injection of oligomycin and inhibition after treatment with BAY-876 (10-0.625µM) in MDA-MB-231 cells. **F.** 4T1 extracellular acidification rates (ECAR (mpH/min)) vs. time after injection of i) treatment or DMSO ii) glucose iii) oligomycin and iv) 2-deoxyglucose. **G.** calculated ECAR after addition of glucose which indicates extent of cellular glycolysis and inhibition from treatment of BAY-876 (10-0.625µM) in 4T1 cells. **H.**: ECAR indicating glycolytic capacity after uncoupling mitochondrial ATP synthesis after injection of oligomycin and inhibition after treatment with BAY-876 (10-0.625µM) in 4T1 cells. **D, E, G, H:** results are representative of 3 biological replicates experiments ± SEM. Statistical analysis was carried out via Dunnett’s one-way ANOVA compared to DMSO control (*p<0.05, **p<0.01, ***p<0.001, ****p<0.0001).

### GLUT1 Inhibitor BAY-876 inhibits glycolysis to impinge on cancer cell proliferation

BAY-876 is an experimental GLUT1 inhibitor that inhibits glycolysis by blocking the uptake of glucose into the cell.^50^ In this regard, we hypothesized that the inhibition of glycolysis by BAY-876 should impinge on cancer cell proliferation. To test this, 3-(4,5-dimethylthiazol-2-yl)-2,5-diphenyltetrazolium bromide (MTT, reporting on cellular redox status and metabolic activity) and sulforhodamine B (SRB, reporting on total cellular protein content) colorimetric assays were performed in cells treated with varying concentrations of BAY-876. Based on potential effects of metabolism inhibition on mitochondrial function with test compounds, SRB assays were performed to corroborate results found in mitochondrial-based MTT assays. Interestingly, in the more glycolytic MDA-MB-231 cells, BAY-876 was found to have EC_50_ values of >100 and 20.47±8.89 µM assessed via MTT and SRB respectively. When compared to the more oxidative cell line 4T1, where BAY-876 was found to have EC_50_ values of 0.21±0.02 and 0.13±0.04 µM assessed via MTT and SRB respectively (**Table 1**). The substantial increase in potency of BAY-876 in 4T1 when compared to MDA-MB-231 cells could be attributed to a large flux of glucose required to fuel mitochondrial OxPhos as the primary ATP source in 4T1 cells (Fig 1A). Associated with the metabolic rates exhibited by 4T1 cells, they also have increased growth rates when compared to MDA-MB-231 (**Fig. 1B**), which may make them more sensitive to inhibition of glucose import.

To further examine the glycolytic effects of BAY-876 on the tested cell lines, we employed Seahorse glycolysis stress tests that report on glycolytic rates as a function of extracellular acidification (ECAR) from end-product lactic acid. Here, glycolysis stress tests on MDA-MB-231 cells illustrated a dose-dependent decrease in glycolytic ECAR from 10-0.625 µM (**Fig. 1C&D**). Additionally, glycolytic capacity was decreased across the treatment concentrations (**Fig. 1C&E**). Intriguingly, glycolysis stress test on 4T1 cells showed that after BAY-876 treatment from 10-0.625 µM, there was a comparably more drastic suppression of glycolytic ECAR when compared to MDA-MB-231 (**Fig. 1F&G**). Glycolytic capacity was also reduced in 4T1 after treatment with BAY-876 (**Fig. 1F&H**). This data supports the notion that the extent by which GLUT1 inhibiting BAY-876 results in functional decreases in glycolysis may be regulated by the metabolic state of the cell. Here, although the ratio of glycolysis is higher in MDA-MB-231 cells, the combined magnitude of glycolysis and respiration is higher in 4T1 cells (**Fig. 1A**) where higher metabolic flux results in an enhanced potency of BAY-876 toward glycolysis (**Fig. 1F-H**) and downstream cellular survival and proliferation (**Table 1**). Acute changes in OCR were not observed in either cell line after treatment with BAY-876 in glycolysis stress tests, indicating that GLUT1 inhibition was not sufficient to impinge on mitochondrial respiration and thus, detailed mitochondrial stress tests were not employed (**Fig. S1**).

### Clinically used Niclosamide and IMD-0354 exhibit mitochondrial uncoupling properties and inhibit cancer cell proliferation

To further expand the pharmacological applications of metabolism targeting agents, we identified two suitable and clinically used candidates: niclosamide and IMD-0354. Niclosamide, an anthelmintic agent with well documented mitochondrial uncoupling properties^51–53^, and IMD-0304, a structural homolog of niclosamide clinically used for atopic dermatitis were selected.^54^ Based on their structural similarity, known mitochondrial targeting capacity, and clinical utility, we further investigated these compounds for their ability to inhibit mitochondrial respiration and cancer cell proliferation in our system. Here, niclosamide was found to inhibit cancer cell proliferation with EC_50_ values against MDA-MB-231 of 4.67±0.99 and 0.87±0.11 µM assessed via MTT and SRB, respectively (**Table 1**). Against 4T1, niclosamide exhibited enhanced potency with EC_50_ values of 0.30±0.06 and 0.36±0.05 µM assessed via MTT and SRB, respectively (**Table 1**). To validate mitochondrial targeting capacity and corroborate cancer cell proliferation studies, we performed SeahorseXF-based mitochondrial stress tests at varying concentrations of niclosamide (**Fig. 2**). In MDA-MB-231, niclosamide injection at 1-0.0625 µM caused an acute increase in OCR (**Fig. 2A**) and a simultaneous increase in ECAR indicating an increase in compensatory glycolysis (**Fig. S2A&B**), consistent with an acute mitochondrial uncoupling effect similar to that of widely used chemical uncouplers such as FCCP. After the addition of ATP-synthase inhibitor oligomycin, there was an apparent biphasic hormetic proton leak response across the test concentrations, with 0.25 µM inducing the largest proton leak (**Fig. 2B**). The %ΔECAR after oligomycin injection was found to be elevated in control groups which indicates a shift to glycolysis after loss of mitochondrial ATP synthesis (**Fig. S2C**). In niclosamide treated groups, the %ΔECAR after oligomycin injection was near zero, indicating that these groups were already uncoupled and insensitive to oligomycin induced changes to ECAR (**Fig. S2C**). Hormetic proton leak responses are consistent with the notion that mitochondrial uncouplers work at an optimal and intermediate concentration to promote maximal respiration, and in this case, concomitant proton leak after oligomycin injection is observed at intermediate concentration ranges (**Fig. 2B**). Additionally, FCCP-induced maximal respiration was decreased in a dose-dependent fashion (**Fig. 2C**) and spare respiratory capacity was significantly reduced across all concentrations, again consistent with these cells already reaching maximal respiration from niclosamide treatment (**Fig. 2D**). The mitochondrial stress test on 4T1 cells illustrated similar metabolic responses when compared to MDA-MB-231, where niclosamide injection at 1-0.0625 µM caused an acute increase in OCR (**Fig. 2E**) and a simultaneous acute increase in ECAR indicating a shift to compensatory glycolysis (**Fig. S2D&E**). After injection of oligomycin there was again a biphasic hormetic proton leak response in the treated groups with 0.5, 0.25, and 0.125 µM causing a maximum proton leak (**Fig. 2F**). In niclosamide treated groups, the %ΔECAR after oligomycin injection was near zero, indicating that similarly to MDA-MB-231, these groups were already uncoupled and insensitive to oligomycin induced changes to ECAR (**Fig. S2F**). FCCP stimulated maximal respiration was reduced in a dose-dependent fashion, with 0.0625 µM alone having an elevated response to FCCP (**Fig. 2G**). As observed in MDA-MB-231, there was a significant loss of spare respiratory capacity across all niclosamide treatment concentrations in 4T1 (**Fig. 2H**). Taken together, the mitochondrial stress test illustrates that niclosamide treatment results in mitochondrial dysfunction with acute oxygen consumption patterns mimicking that of a mitochondrial uncoupling agent. Mitochondrial perturbations are accompanied by acute increases in extracellular acidification rates that can be attributed to acute compensatory glycolysis, further highlighting the ability of cancer cells to upregulate metabolic pathways to sustain bioenergetic demands.

**Figure 2:**
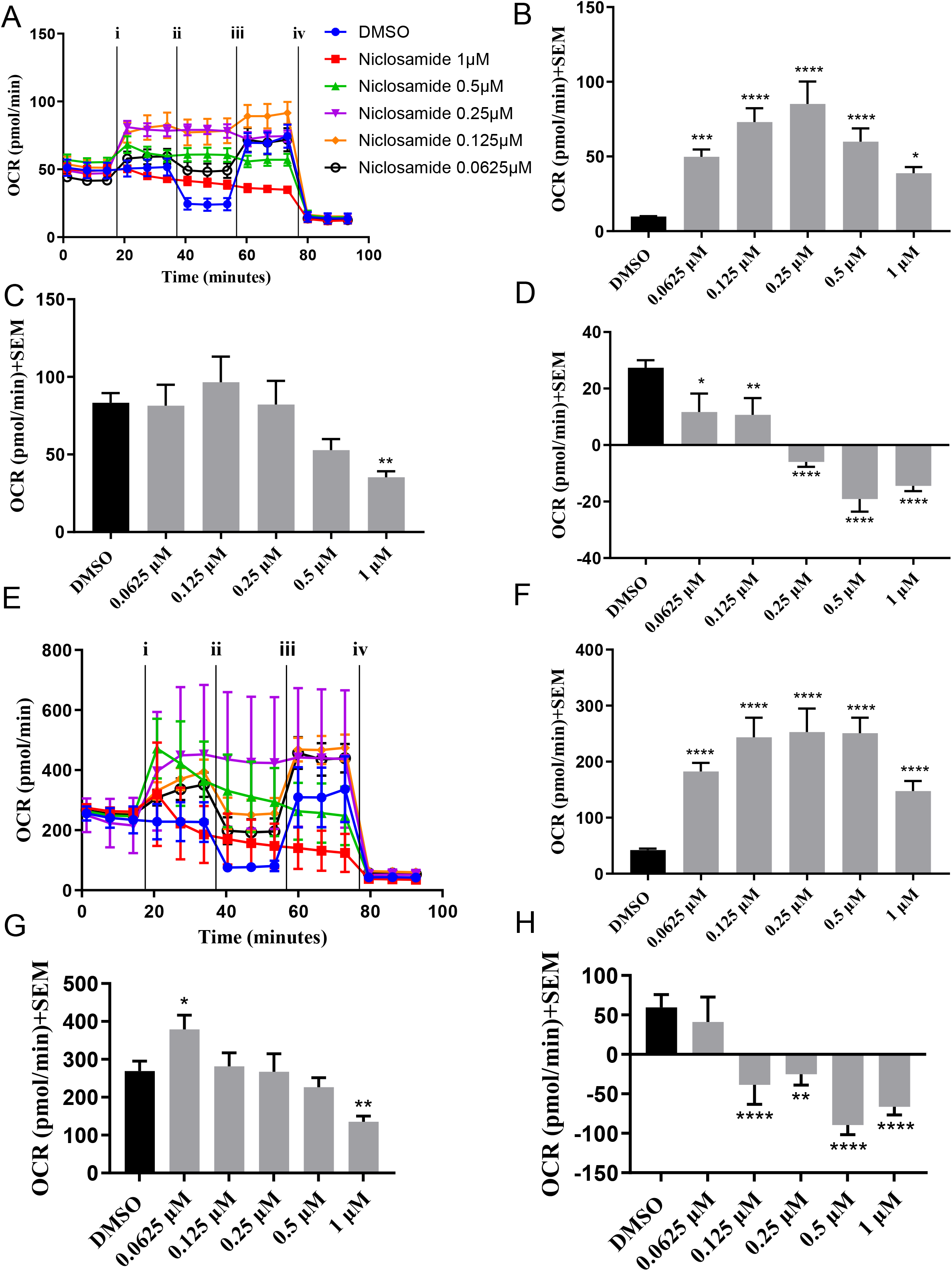
**A.** MDA-MB-231 oxygen consumption rates (OCR (pmol/min)) vs. time after injection of i) treatment or DMSO ii) oligomycin iii) FCCP and iv) Rotenone and Antimycin A after treatment with niclosamide (1-0.0625µM). **B.** calculated OCR subtracting non-mitochondrial respiration from OCR after oligomycin of DMSO control and after acute niclosamide injection (1-0.0625µM) indicating proton leak in MDA-MB-231 cells. **C.** Maximal respiration OCR of DMSO control and after treatment with niclosamide (1-0.0625µM) in MDA-MB-231 cells. **D.** Spare respiratory capacity OCR from subtracting OCR before oligomycin injection from maximal respiration after treatment with niclosamide (1-0.0625µM) in MDA-MB-231 cells. **E.** 4T1 oxygen consumption rates (OCR (pmol/min)) vs. time after injection of i) treatment or DMSO ii) oligomycin iii) FCCP and iv) Rotenone and Antimycin A after treatment with niclosamide (1-0.0625µM). **F.** calculated OCR subtracting non-mitochondrial respiration from OCR after oligomycin of DMSO control and after acute niclosamide injection (1-0.0625µM) indicating proton leak in 4T1 cells. **G.** Maximal respiration OCR of DMSO control and after treatment with niclosamide (1-0.0625µM) in 4T1 cells. **H.** Spare respiratory capacity OCR from subtracting OCR before oligomycin injection from maximal respiration after treatment with niclosamide (1-0.0625µM) in 4T1 cells. **B, C, D, F, G, H**: results are representative of 3 biological replicates experiments ± SEM. Statistical analysis was carried out via Dunnett’s one-way ANOVA compared to DMSO control (*p<0.05, **p<0.01, ***p<0.001, ****p<0.0001).

The chemical properties of IMD-0354 largely mimic that of niclosamide, wherein the phenolic moiety likely exhibits similar uncoupling activity. In a similar fashion to niclosamide, we evaluated the cell proliferation inhibition properties, along with the mitochondrial and glycolysis stress tests of IMD-0354. Here, IMD-0354 was found to have EC_50_ values of cell proliferation against MDA-MB-231 of 3.43±0.97 and 0.75±0.09 µM assessed via MTT and SRB respectively (**Table 1**). Against 4T1, IMD-0354 was found to have EC_50_ values of 0.27±0.1 and 0.26±0.09 µM assessed via MTT and SRB respectively (**Table 1**). In a mitochondrial stress test on MDA-MB-231, IMD-0354 treatment at a concentration range of 1.25-0.078 µM saw an acute increase in OCR (**Fig. 3A**) and a biphasic hormetic acute increase in ECAR (**Fig. S3A&B**). Similarly, after oligomycin injection, there was again a biphasic hormetic proton leak in IMD-0354 treated groups (**Fig. 3B**). This response mirrored that of the acute ECAR hormetic response, with 0.3125 µM giving the largest increase in ECAR and proton leak respectively (**Fig. 3B**). There was a nearly zero %ΔECAR in IMD-0354 after oligomycin treatment, indicating no further increase to glycolytic metabolism and as a result, an apparent oligomycin insensitivity (**Fig. S3C**). Additionally, there was a dose-dependent decrease in FCCP stimulated maximal respiration and spare respiratory capacity (**Fig. 3C&D**), again likely due to the mitochondria functioning at maximal respiration, where an additional load of uncoupling agent (FCCP) results in reduced maximal OCR. The mitochondrial stress test on 4T1 cells illustrated an acute increase in OCR after injection of IMD-0354 (**Fig. 3E**). Like MDA-MB-231, in 4T1 cells there was a hormetic acute increase in ECAR (**Fig. S3D&E**) and hormetic proton leak after oligomycin injection (**Fig. 3F**), with 0.3125 µM also giving rise to the largest response for both. Additionally, IMD-0354 increased maximal respiration at lower concentrations, and decreased at higher concentrations (**Fig. 3G**), results that extended to decreases in spare respiratory capacity (**Fig. 3H**). There was a dose-dependent decrease in %ΔECAR after oligomycin (**Fig. S3F**) indicating that there was no further stimulation of compensatory glycolysis and that cells are insensitive to oligomycin. As observed with niclosamide, chemically similar IMD-0354 exhibited metabolic effects consistent with mitochondrial uncoupling capacity.

**Figure 3:**
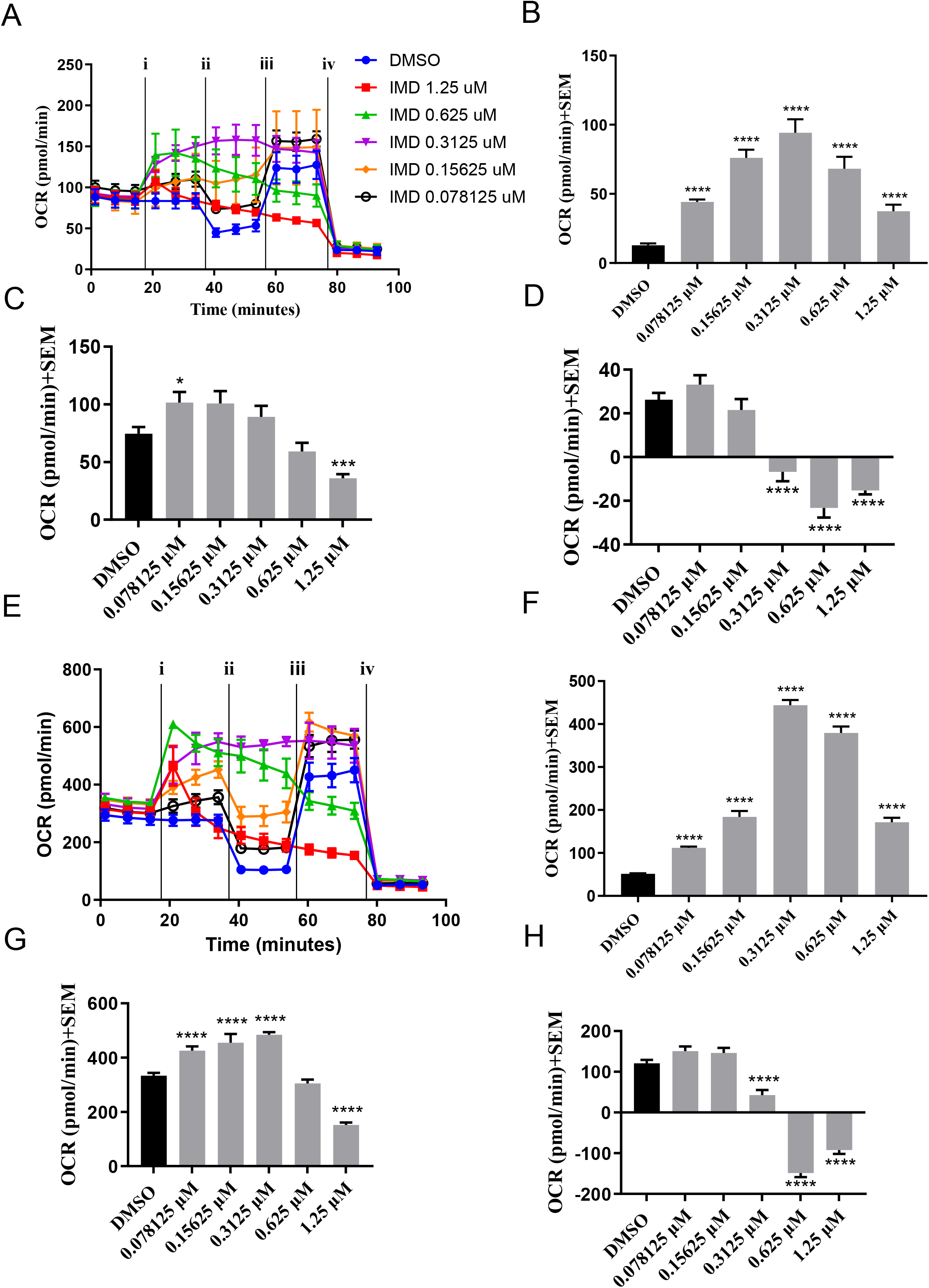
**A.** MDA-MB-231 oxygen consumption rates (OCR (pmol/min)) vs. time after injection of i) treatment or DMSO ii) oligomycin iii) FCCP and iv) Rotenone and Antimycin A after treatment with IMD-0354 (1.25-0.078125µM). **B.** Calculated OCR subtracting non-mitochondrial respiration from OCR after oligomycin of DMSO control and after acute IMD-0354 injection (1.25-0.078125µM) indicating proton leak in MDA-MB-231 cells. **C.** Maximal respiration OCR of DMSO control and after treatment with IMD-0354 (1.25-0.078125µM) in MDA-MB-231 cells. **D.** Spare respiratory capacity OCR from subtracting OCR before oligomycin injection from maximal respiration after treatment with IMD-0354 (1.25-0.078125µM) in MDA-MB-231 cells. **E.** 4T1 oxygen consumption rates (OCR (pmol/min)) vs. time after injection of i) treatment or DMSO ii) oligomycin iii) FCCP and iv) Rotenone and Antimycin A after treatment with IMD-0354 (1.25-0.078125µM). **F.** Calculated OCR subtracting non-mitochondrial respiration from OCR after oligomycin of DMSO control and after acute IMD-0354 injection (1.25-0.078125µM) indicating proton leak in 4T1 cells. **G.** Maximal respiration OCR of DMSO control and after treatment with IMD-0354 (1.25-0.078125µM) in 4T1 cells. **H.** Spare respiratory capacity OCR from subtracting OCR before oligomycin injection from maximal respiration after treatment with IMD-0354 (1.25-0.078125µM) in 4T1 cells. **B, C, D, F, G, H:** results are representative of 3 biological replicates experiments ± SEM. Statistical analysis was carried out via Dunnett’s one-way ANOVA compared to DMSO control (*p<0.05, **p<0.01, ***p<0.001, ****p<0.0001).

### Pyrvinium pamoate (PP) targets mitochondrial metabolism and inhibits cancer cell proliferation

A cyanine-based compound, PP is a delocalized lipophilic cation (DLC), and therefore it is characteristically expected to influence mitochondria due to a higher accumulation in the mitochondrial matrix.^16,17^ PPs cytotoxic effects have been shown to be under nutrient restriction conditions highlighting the potential utility of this anthelmintic agent to exhibit mitochondrial targeting anticancer properties, qualifying PP to be evaluated for its antiproliferative and metabolic properties in our system. PP was found to have EC_50_ values against MDA-MB-231 of 0.49±0.05 and 0.15±0.00 µM assessed via MTT and SRB respectively (**Table 1**). Against 4T1, PP was found to have EC_50_ values of 0.034±0.024 and 0.020±0.01 µM assessed via MTT and SRB respectively (**Table 1**). In a mitochondrial stress test on MDA-MB-231, PP caused an acute decrease in OCR (**Fig. 4A&B**) and a dose-dependent increase in acute ECAR (**Fig. S4A&B**), indicating that mitochondrial respiration was being inhibited while a dose-dependent switch to a compensatory glycolytic metabolism was likely occurring. After oligomycin injection, PP treatment led to a dose-dependent decrease in OCR associated with mitochondrial ATP production (**Fig. 4C**). In addition, there was a dose-dependent sensitivity to oligomycin as evidenced by a concomitant decrease in %ΔECAR after oligomycin injection (**Fig. S4C**). FCCP stimulated maximal respiration was found to be reduced at higher concentrations of PP (**Fig. 4D**, 1 and 0.5 µM) and spare respiratory remained consistent with DMSO control aside from 0.125 and 0.25 µM which saw an increase in associated OCR (**Fig. 4E**), potentially indicating a tightly controlled dose-dependency with regard to the ability of mitochondria to respond to additional metabolic stress in the presence of PP. This is unlike what was observed with protonophore’s niclosamide and IMD-0354, where PP remained sensitive to stimulation by FCCP as evidenced by the retention or increase in spare respiratory capacity (**Fig. 4E**). This suggests that PP does not interfere with mitochondria through perturbations in the proton gradient, which is consistent with the structure of PP lacking labile proton moieties present in known uncoupling agents (ex. FCCP, dinitrophenol, niclosamide, etc). In the mitochondrial stress test on 4T1 cells, there was an acute decrease in OCR across treatment groups (**Fig. 4F&G**). Additionally, there was a simultaneous increase in acute ECAR indicating a switch to compensatory glycolysis (**Fig. S4D&E**). Interestingly, the increase in acute ECAR in 4T1 cells did not occur in a dose-dependent fashion as seen with MDA-MB-231. This could reflect a greater need for glycolytic compensation in 4T1 cells as most of their ATP production occur through OxPhos. The increase in ECAR even at lower concentrations of PP could be due to a higher mitochondrial content in 4T1 causing a larger accumulation of PP, giving rise to a higher potency of PP to induce compensatory glycolysis. After oligomycin injection, there was a PP dose-dependent decrease in OCR associated with mitochondrial ATP production (**Fig. 4H**), but cells compensatory glycolysis was completely insensitive to oligomycin with near zero %ΔECAR (**Fig. S4F**). Additionally, there was a dose-dependent decrease in FCCP stimulated maximal respiration to a greater extent in 4T1 cells **(Fig. 4I**) than in MDA-MB-231 cells, but spare respiratory capacity was retained (**Fig. 4J**). These results indicate that PP exhibits acute and potent mitochondrial targeting capacity, and in a mechanistically different fashion compared to mitochondrial respiration profiles of cells treated with niclosamide or IMD-0354. These results substantiate the hypothesis that the delocalized lipophilic cation structure of PP promotes its mitochondrial accumulation, but specific target is still unknown. However, compensatory glycolysis upon mitochondrial dysfunction with PP is consistent with results obtained with uncoupling agents and qualifies glycolysis inhibitors as a potential combination strategy.

**Figure 4:**
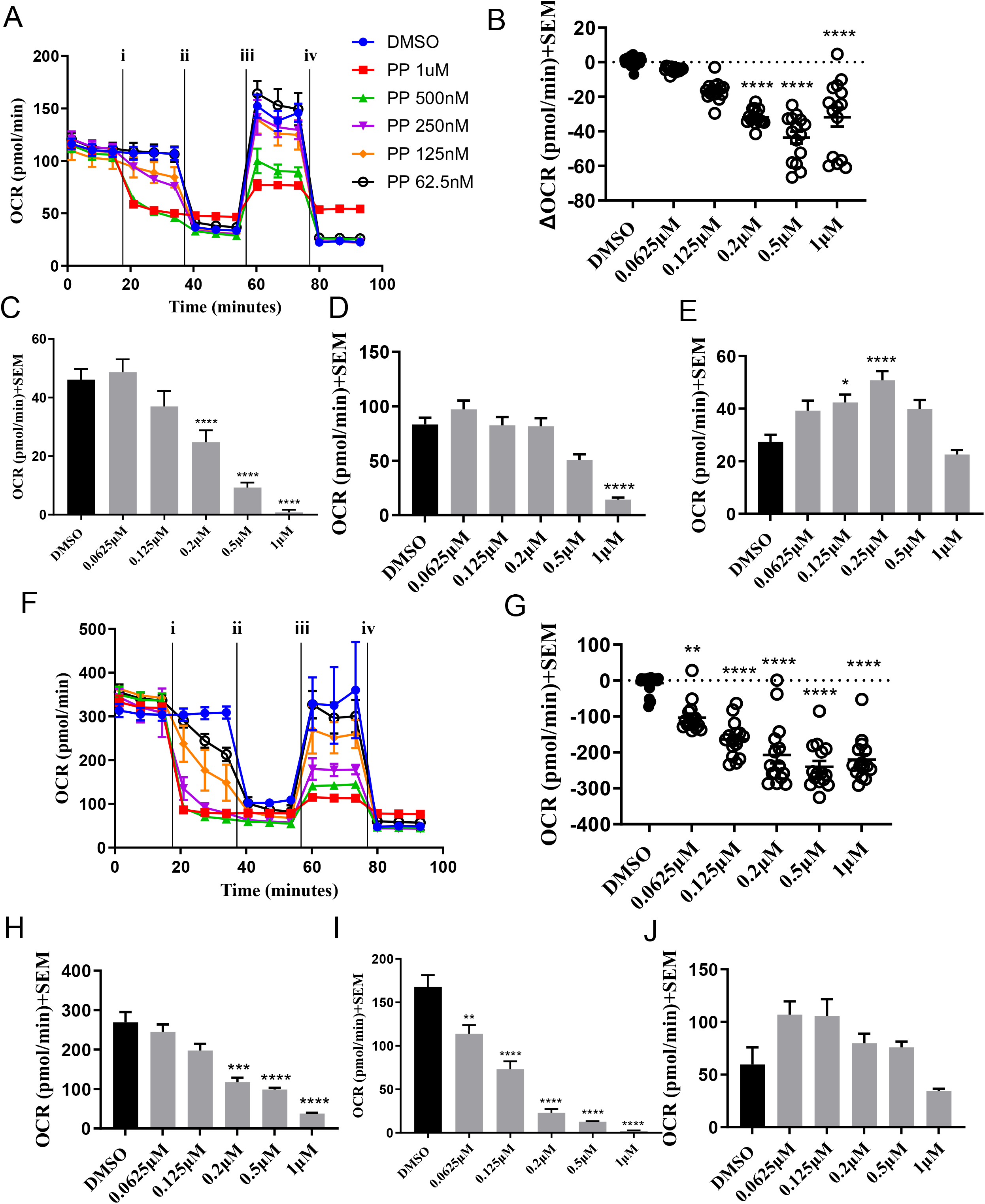
**A.** MDA-MB-231 oxygen consumption rates (OCR (pmol/min)) vs. time after injection of i) treatment or DMSO ii) oligomycin iii) FCCP and iv) Rotenone and Antimycin A after treatment with PP (1-0.0625µM). **B.** ΔOCR after acute injection of PP (1-0.0625µM) in MDA0-MB-231 cells. **C.** ΔOCR subtracting OCR before injection of FCCP from before oligomycin injection of DMSO control and after acute PP injection (1-0.0625µM) indicating mitochondrial ATP production in MDA-MB-231 cells. **D.** Maximal respiration OCR of DMSO control and after treatment with PP (1-0.0625µM) in MDA-MB-231 cells. **E.** Spare respiratory capacity OCR from subtracting OCR before oligomycin injection from maximal respiration after treatment with PP (1-0.0625µM) in MDA-MB-231 cells. **F.** 4T1 oxygen consumption rates (OCR (pmol/min)) vs. time after injection of i) treatment or DMSO ii) oligomycin iii) FCCP and iv) Rotenone and Antimycin A after treatment with PP (1-0.0625µM). **G.** ΔOCR after acute injection of PP (1-0.0625µM) in MDA0-MB-231 cells. **H.** ΔOCR subtracting OCR before injection of FCCP from before oligomycin injection of DMSO control and after acute PP injection (1-0.0625µM) indicating mitochondrial ATP production in 4T1 cells. **I.** Maximal respiration OCR of DMSO control and after treatment with PP (1-0.0625µM) in 4T1 cells. **J.** Spare respiratory capacity OCR from subtracting OCR before oligomycin injection from maximal respiration after treatment with PP (1-0.0625µM) in 4T1 cells.

### Combination strategies targeting glycolysis and mitochondrial metabolism result in metabolic crisis in metabolically variable 4T1 and MDA-MB-231 breast cancer cells

Our results indicate that inhibition of mitochondrial metabolism with niclosamide, IMD-0354, or PP results in an acute increase in complementary glycolysis, respectively. Cancer cells characteristically exhibit the ability to alter their metabolic pathways to compensate for defects (ex. pharmacological inhibition, mutations in metabolic genes, nutrient deprivation/oxygen gradients) and hence can overcome single agent treatment with glycolysis inhibitors or mitochondrial targeting agents.^46–49^ In this regard, we sought to combine GLUT1 inhibitor BAY-876 with mitochondrial targeting agents niclosamide, IMD-0354, and PP. The mitochondrial stress test on MDA-MB-231 after treatment with a combination of BAY-876+niclosamide (B/N) at 1-0.0625 µM resulted in acute increases in OCR which was maintained for the duration of the experiment (**Fig 5A**). There was a simultaneous spike in ECAR in a dose-dependent fashion after acute B/N injection that reduced over time (**Fig. 5B**). This suggests that niclosamide initially increases compensatory glycolysis but over time BAY-876 inhibits GLUT1 and thus compensatory glycolysis decreases due to lack of glucose. As seen with IMD-0354 as a single agent, following oligomycin injection there was a biphasic hormetic proton leak after B/N treatment with 0.25 µM achieving the largest proton leak (**Fig. 5C**). There was a near zero %ΔECAR after oligomycin indicating that no further compensatory glycolysis was induced, and cells were insensitive to oligomycin (**Fig. 5B**). There was a hormetic response to FCCP which induced maximal respiration with 0.25 µM of B/N inducing the largest increase in FCCP induced maximal respiration (**Fig. 5D**), however spare respiratory capacity was significantly reduced for all concentrations of B/N (**Fig. 5E**).

**Figure 5:**
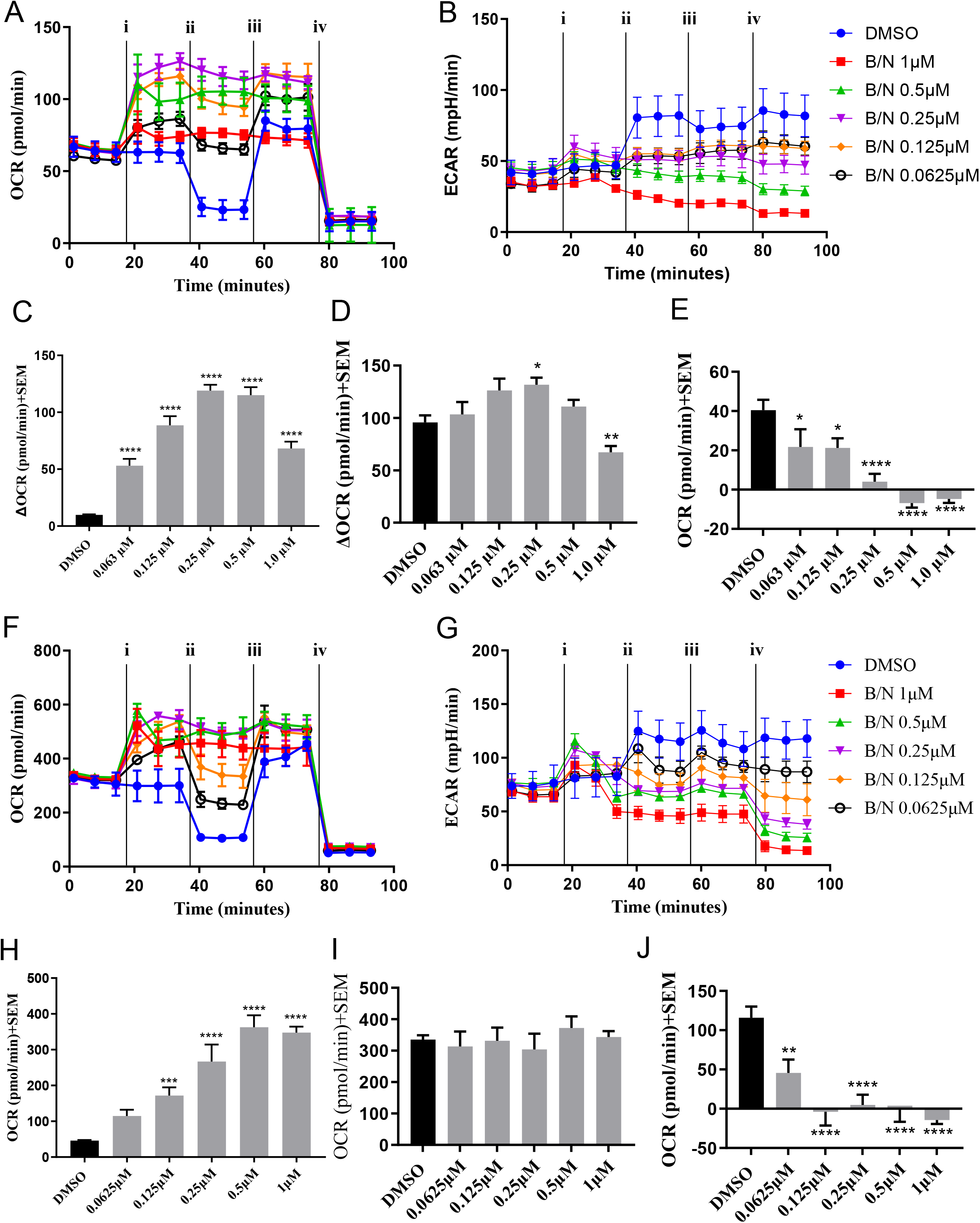
**A.** MDA-MB-231 oxygen consumption rates (OCR (pmol/min)) vs. time after injection of i) treatment or DMSO ii) oligomycin iii) FCCP and iv) Rotenone and Antimycin A after treatment with B/N (1-0.0625 µM). **B.** MDA-MB-231 extracellular acidification rates (ECAR (mpH/min)) vs. time after injection of i) treatment or DMSO ii) oligomycin iii) FCCP and iv) Rotenone and Antimycin A after treatment with B/N (1-0.0625 µM). **C.** ΔOCR subtracting non-mitochondrial respiration from OCR after oligomycin of DMSO control and after acute B/N injection (1-0.0625 µM) indicating proton leak in MDA-MB-231 cells. **D.** Maximal respiration OCR of DMSO control and after treatment with B/N (1-0.0625 µM) in MDA-MB-231 cells. **E**. Spare respiratory capacity OCR from subtracting OCR before oligomycin injection from maximal respiration in MDA-MB-231 cells. **F.** 4T1 oxygen consumption rates (OCR (pmol/min)) vs. time after injection of i) treatment or DMSO ii) oligomycin iii) FCCP and iv) Rotenone and Antimycin A after treatment with B/N (1-0.0625 µM). **G.** 4T1 extracellular acidification rates (ECAR (mpH/min)) vs. time after injection of i) treatment or DMSO ii) oligomycin iii) FCCP and iv) Rotenone and Antimycin A after treatment with B/N (1-0.0625 µM). **H.** ΔOCR subtracting non-mitochondrial respiration from OCR after oligomycin of DMSO control and after acute B/N injection (1-0.0625 µM) indicating proton leak in 4T1 cells. **I.** Maximal respiration OCR of DMSO control and after treatment with B/N (1-0.0625 µM) in 4T1 cells. **J**. Spare respiratory capacity OCR from subtracting OCR before oligomycin injection from maximal respiration in 4T1 cells. **C, D, E, H, I, J:** results are representative of 3 biological replicates experiments ± SEM. Statistical analysis was carried out via Dunnett’s one-way ANOVA compared to DMSO control (*p<0.05, **p<0.01, ***p<0.001, ****p<0.0001).

In the mitochondrial stress test on 4T1 cells with treatment of B/N there was an increase in OCR after acute injection of B/N (**Fig. 5F**). Additionally, there was an acute spike in ECAR which reduced over time in a dose-dependent fashion indicating an initial switch to compensatory glycolysis followed by time dependent inhibition by BAY-876 (**Fig. 5G**). No further stimulation of glycolysis was found after oligomycin injection as evidenced by a near zero %ΔECAR, indicating that GLUT1-inhibited cells were insensitive to oligomycin. As a single agent, niclosamide induced a hormetic proton leak (**Fig. 2F**). In combination with BAY-876, there was a hormetic induced proton leak but the addition of BAY-876 decreased sensitivity shifting the hormetic curve toward increased concentrations (**Fig. 5H**). This result may suggest that the ability of niclosamide to affect mitochondrial metabolism may depend to some extent on glycolytic metabolites (ex. pyruvate), or glycolysis-derived reducing equivalents (ex. NAD(P)H). Also, it is possible that the inhibited glucose flux with BAY-876 alters mitochondrial activity such that the effective uncoupling concentration of niclosamide is increased, highlighting the dynamic interplay between glycolysis and OxPhos that can dictate drug sensitivity. There was no effect on FCCP induced maximal respiration when compared to control after treatment with B/N (**Fig. 5I**), but spare respiratory capacity was reduced across all treatment concentrations (**Fig. 5G**).

The mitochondrial stress test on MDA-MB-231 showed that after treatment with BAY-876+pyrvinium pamoate (B/PP) at 1-0.0625 µM caused a decrease in OCR after injection (**Fig. 6A**). Simultaneously, there was a dose-dependent increase in ECAR which reduced over time indicating that the initial compensatory glycolysis was being inhibited over time from lack of glucose due to BAY-876 inhibition of GLUT1 (**Fig. 6B**), similar to what was observed with B/N treatment. After oligomycin there was a dose-dependent decrease in proton leak and mitochondrial ATP linked OCR, indicating that mitochondria were not successfully transferring electrons to oxygen but retaining their proton gradient (**Fig. 6C**). Additionally, a dose-dependent decrease in ECAR after oligomycin was observed, indicating that sensitivity to both oligomycin and GLUT1 inhibition is a dose-dependent process (**Fig 6B**). In B/PP treated groups, there was a dose-dependent decrease in both FCCP mediated maximal respiration and spare respiratory capacity (**Fig. 6D-E**).

**Figure 6:**
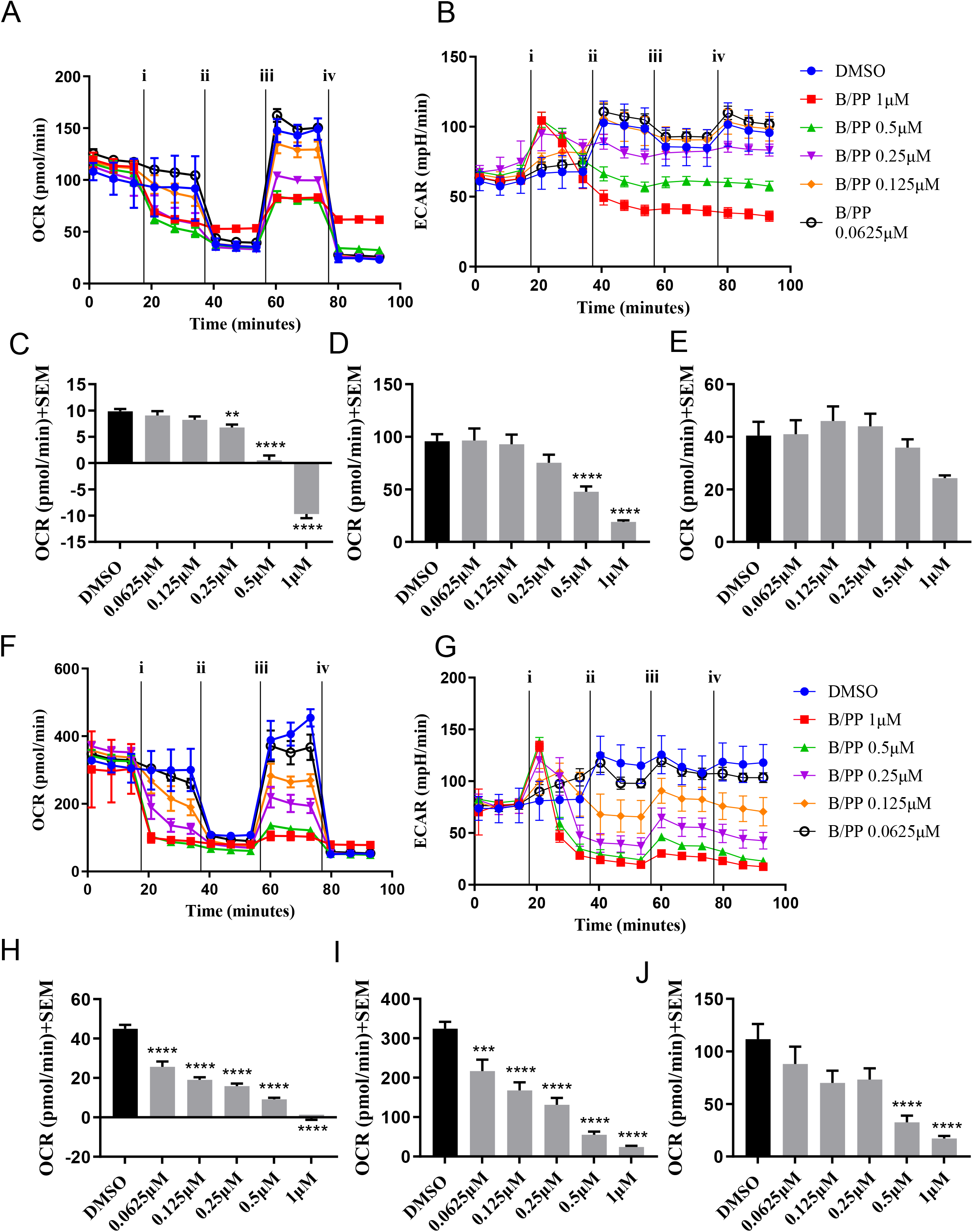
**A.** MDA-MB-231 oxygen consumption rates (OCR (pmol/min)) vs. time after injection of i) treatment or DMSO ii) oligomycin iii) FCCP and iv) Rotenone and Antimycin A after treatment with B/PP (1-0.0625 µM). **B.** MDA-MB-231 extracellular acidification rates (ECAR (mpH/min)) vs. time after injection of i) treatment or DMSO ii) oligomycin iii) FCCP and iv) Rotenone and Antimycin A after treatment with B/PP (1-0.0625 µM). **C.** ΔOCR subtracting non-mitochondrial respiration from OCR after oligomycin of DMSO control and after acute B/PP injection (1-0.0625 µM) indicating proton leak in MDA-MB-231 cells. **D.** Maximal respiration OCR of DMSO control and after treatment with B/PP (1-0.0625 µM) in MDA-MB-231 cells. **E**. Spare respiratory capacity OCR from subtracting OCR before oligomycin injection from maximal respiration in MDA-MB-231 cells. **F.** 4T1 oxygen consumption rates (OCR (pmol/min)) vs. time after injection of i) treatment or DMSO ii) oligomycin iii) FCCP and iv) Rotenone and Antimycin A after treatment with B/PP (1-0.0625 µM). **G.** 4T1 extracellular acidification rates (ECAR (mpH/min)) vs. time after injection of i) treatment or DMSO ii) oligomycin iii) FCCP and iv) Rotenone and Antimycin A after treatment with B/PP (1-0.0625 µM). **H.** ΔOCR subtracting non-mitochondrial respiration from OCR after oligomycin of DMSO control and after acute B/PP injection (1-0.0625 µM) indicating proton leak in 4T1 cells. **I.** Maximal respiration OCR of DMSO control and after treatment with B/PP (1-0.0625 µM) in 4T1 cells. **J**. Spare respiratory capacity OCR from subtracting OCR before oligomycin injection from maximal respiration in 4T1 cells. **C, D, E, H, I, J:** results are representative of 3 biological replicates experiments ± SEM. Statistical analysis was carried out via Dunnett’s one-way ANOVA compared to DMSO control (*p<0.05, **p<0.01, ***p<0.001, ****p<0.0001).

The mitochondrial stress test in 4T1 cells with B/PP treatment showed that there was a decrease in OCR after acute injection across all concentrations (**Fig. 6F**). As seen in MDA-MB-231, there was a simultaneous dose-dependent increase in ECAR after acute injection of B/PP that turned into a decrease over time (**Fig. 6G**). Compensatory glycolysis remained insensitive to oligomycin at all concentrations as evidenced by near or below zero %ΔECAR values (**Fig. 6G**). However, after oligomycin injection, it was found that B/PP treatment resulted in a dose-dependent decrease in proton leak and mitochondrial ATP associated OCR (**Fig. 6H**). Treatment with B/PP caused a dose-dependent decrease in FCCP stimulated maximal respiration and spare respiratory capacity (**Fig. 6I-J**).

In this set of experiments, we sought to understand if by combining agents that inhibit glycolysis or mitochondrial metabolism independently could block the resulting compensatory metabolic pathway. By utilizing a GLUT1 inhibitor (BAY-876), compensatory glycolysis upon treatment with mitochondrial uncoupling agent niclosamide, along with delocalized cation PP, is inhibited. By understanding the underlying metabolic fuel preference of a given cell line, metabolism-targeting treatment strategies can be designed to target the preferred metabolic pathway along with the cellular compensatory mechanisms for durable bioenergetic perturbations.

## CONCLUSIONS

Cancer cells exhibit the capacity to alter metabolic phenotypes in the context of nutritional challenge. Numerous single agent approaches targeting mitochondrial metabolism in cancer have failed due to either dose limiting off target toxicities, or lack of efficacy in vivo (ref). To mitigate these clinical challenges, we illustrate the potential utility of repurposing already FDA approved mitochondrial targeting anthelmintic agents, niclosamide and pyrvinium pamoate, to be combined with GLUT1 inhibitor BAY-876 to enhance the inhibitory capacity of the major metabolic phenotypes exhibited by tumors. Specific responses to mitochondrial and glycolysis targeting agents are altered as a function of basal metabolic rates and fuel preference of the cell line of interest, highlighting the potential to cater metabolism targeting treatment regimens based on specific tumor nutrient handling. These studies warrant further investigation into targeting tumor metabolism as a combination treatment regimen that can be tailored by basal and compensatory metabolic phenotypes.

## MATERIALS AND METHODS

### Materials and equipment

BAY-876 and pyrvinium pamoate were purchased from MedChemExpress (Monmouth Junction, NJ). Niclosamide ethanol amine was purchased from Caymen Chemical (Ann Arbor, MI). IMD-0354 was synthesized using 5-chloro-2-hydroxybenzoic acid, 3,5-bis(trifluoromethyl)aniline, and thionyl chloride all purchased from Ambeed (Arlington Heights, IL). The ^1^H- and ^13^C-NMR spectra were plotted on a Bruker Ascend™ 400 spectrometer. High-resolution mass spectra (HRMS) were recorded using a Bruker micrOTOF-Q III ESI mass spectrometer. Absorbance values were measured using BioTek Synergy 2 Multimode Microplate Reader (BioTek Instruments Inc., Winooski, VT, USA). Metabolic experiments were carried out using a Seahorse XFe96 Extracellular Flux Analyzer (Agilent, Santa Clara, California). All statistical analysis was carried out using GraphPad Prism 10 software (GraphPad Software, Boston, MA, USA).

### Synthesis of 5-chloro-2-hydroxybenzoyl chloride

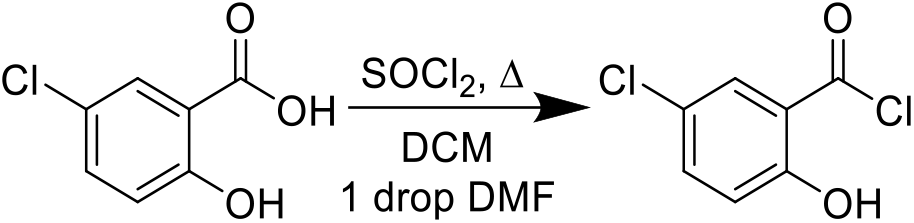

The synthesis of 5-chloro-2-hydroxybenzoyl chloride began by first adding 5-chloro-2-hydroxybenzoic acid (2 mmol) to a round bottom flask with 5mL dichloromethane and a drop of dimethylformamide. To the reaction mixture, thionyl chloride (10 mmol) was added dropwise and the reaction was heated at 50°C for 12 hours. Reaction progress was checked after 12 hours via TLC (70% EtOAc/Hexanes) or for complete dissolution of 5-chloro-2-hydroxybenzoic acid. Upon complete dissolution of the acid, the DCM was removed via rotary evaporation to yield the acid chloride crude. No further purification was carried out and the acid chloride was used within 1 hour of synthesis.

### Synthesis of N-(3,5-bis(trifluoromethyl)phenyl)-5-chloro-2-hydroxybenzamide (IMD-0354)

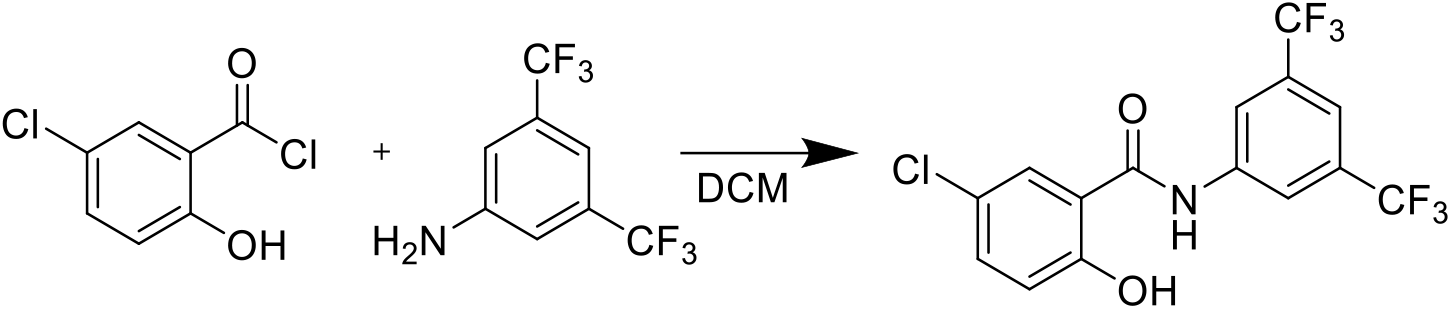

The synthesis of IMD-0354 began by first dissolving 3,5-bis(trifluoromethyl)aniline (2.2 mmol) in 5mL dichloromethane. The previously generated 5-chloro-2-hydroxybenzoyl chloride was then dissolved in 3 mL of dichloromethane and added dropwise to the reaction mixture. After stirring for 1 hour the reaction formed a precipitate which was filtered, rinsed 2x with 3 mL DCM and dried to afford IMD-0354 in 89% yield.

### Spectral characterization of N-(3,5-bis(trifluoromethyl)phenyl)-5-chloro-2-hydroxybenzamide (IMD-0354)

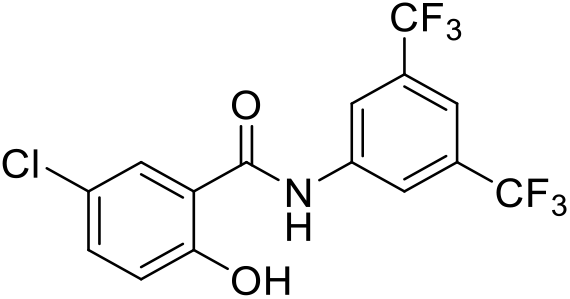

**^1^H NMR (400 MHz, DMSO-d6):** δ 11.46 (s br., 1H), 10.87 (s, 1H), 8.45 (s, 2H), 7.88 (d, 1H, J= 2.6

Hz), 7.83 (s, 1H), 7.49 (dd, 2H, J= 2.6, 8.8 Hz), 7.06 (d, 1H, J=8.8 Hz)

**^13^C NMR (100 MHz, DMSO-d6):** δ 165.99, 156.83, 140.71, 133.71, 131.18 (q, J= 32 Hz), 129.03,

### 123.68 (q, J= 271 Hz), 123.25, 120.80 (d, J=3.6 Hz), 120.51, 119.52, 117.30 (t, J=3.6 Hz)

**^19^F NMR (376 MHz, DMSO-d6)**: δ −61.6602

**HRMS (ESI) m/z:** calculated for C_15_H_8_ClF_6_NO_2_ [M-1]^-^: 382.0064, found 382.0263

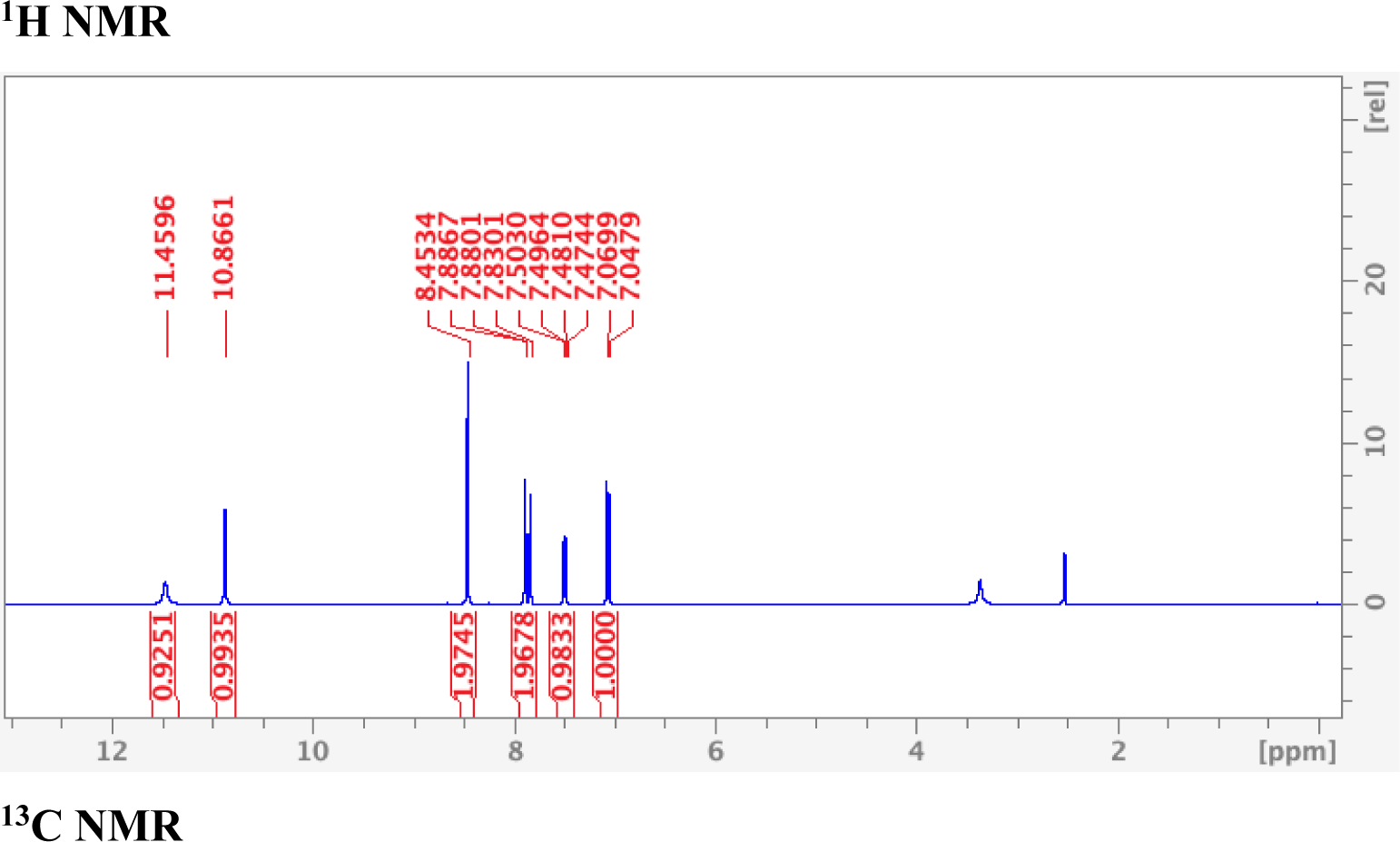
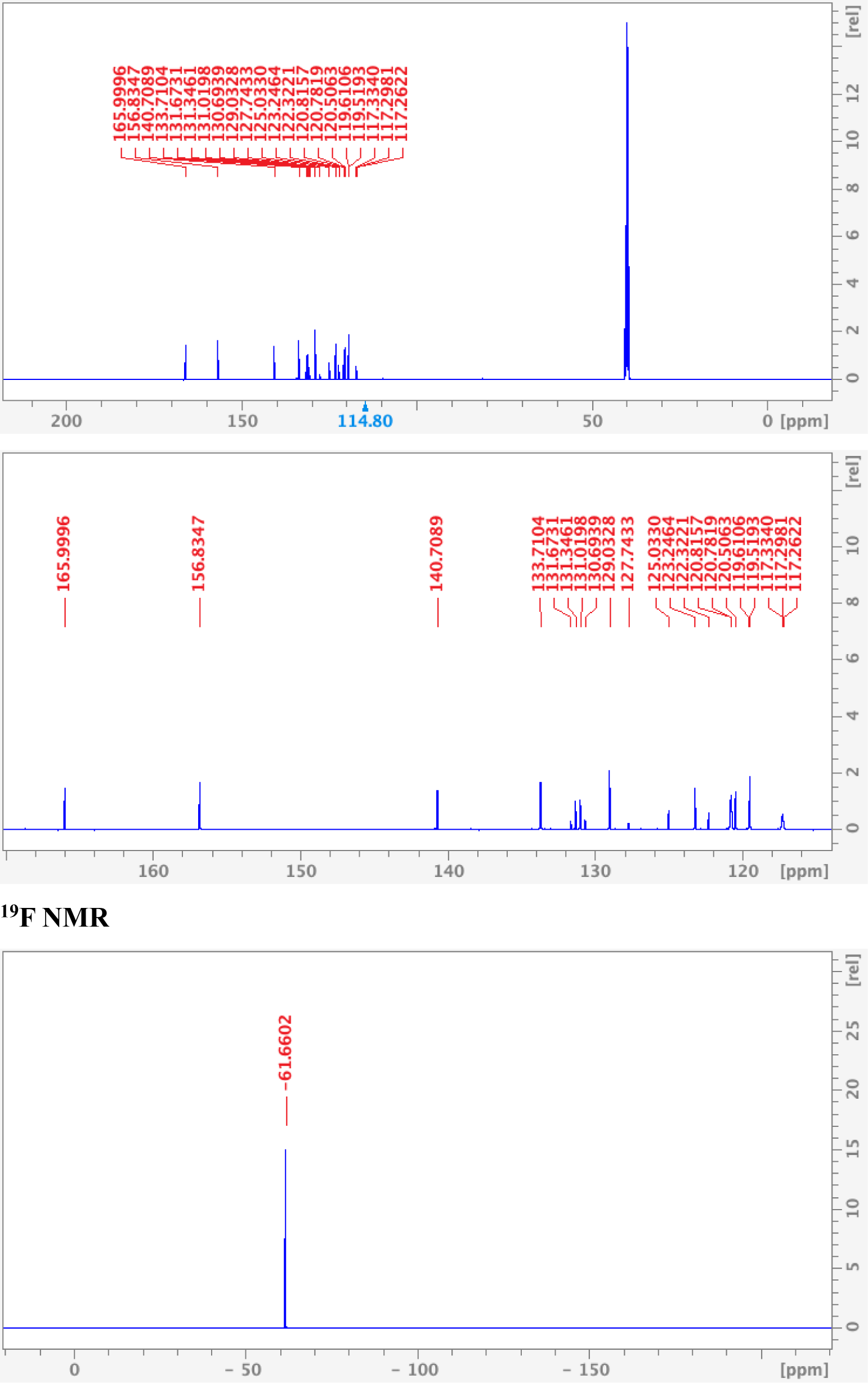

### Cell lines and culture conditions

MDA-MB-231 cells (ATCC) were grown in DMEM supplemented with 10% FBS and penicillin-streptomycin (50 μg/mL). 4T1 cells (ATCC) were cultured in RPMI-1640 supplemented with 10% FBS and penicillin-streptomycin (50 μg/mL). All cells were incubated at 37C and 5% CO_2_.

### MTT Cell Proliferation Assay

Confluent cell cultures were treated with trypsin and resuspended at 5×10^4^ cells/mL. To a 96-well plate, 100 µl of the 5×10^4^ cells/mL solution was added and allowed to incubate at 37°C, 5% CO_2_ for 24 hours. Compounds were then added and allowed to incubate for 3 days. At this time 10 µL of MTT (5 mg/mL) was added to the 96-well plate and further incubated for 4 hours. Following the 4-hour incubation with MTT, 100 µL of SDS (0.1 g/mL, 0.01N HCl) was added and 96-well plates were allowed to incubate for an additional 4 hours. Absorbance values were then taken at 570 nm using a Biotek Synergy 2 plate reader. Treatment absorbances are taken as percentages of the average untreated absorbances and plotted against log(concentration) in GraphPad Prism 10 to generate effective concentration values were 50% of the cells are not proliferating (EC_50_).

### SRB Cell Proliferation Inhibition Assay

Cells were seeded (3×10^4^cells/well) in 48 well plates and incubated overnight (37°C, 5%CO_2_). Test compounds were then exposed to cells in duplicate in serial dilution fashion and were incubated for an additional 72 hours (37°C, 5%CO_2_). Media was then aspirated, and cells were washed three times with cold PBS and left overnight to dry-fix. 100µL Sulforhodamine B (SRB, 0.5% w/v in 1% aqueous acetic acid) was added to each well and was incubated at 37°C for 1hr. SRB was rinsed with 1% acetic acid solution and dried. Dyed cells were then lysed with Tris (10mM, pH 10) and absorbance readings were recorded at 540nm of each well. Treatment absorbances are taken as percentages of the average untreated absorbances and plotted against log(concentration) in GraphPad Prism 10 to generate effective concentration values were 50% of the cells are not proliferating (EC_50_).

### MDA-MB-231 and 4T1 growth curve

MDA-MB-231 and 4T1 cells were seeded at 30,000 cells/well in a 48 well plate. After 24 hours and every subsequent day, media was aspirated from 6 wells, rinsed three times with cold PBS and allowed to dry fix for a minimum of 24 hours. After all wells had dry fixed for 24 hours, 0.5% w/v SRB in 1% acetic acid was added to the wells for 45 minutes, aspirated and rinsed three times with 1% acetic acid, allowed to dry, solubilized in 10mM Tris bas (pH 10.2) and absorbance was measured at 540 nm. Growth curve plot was generated by plotting A_540_ vs day.

### Seahorse XFe96 MitoStress Test

MDA-MB-231 or 4T1 cells were seeded on a Seahorse XFe96 96-well plate at 20,000 cells/well and incubated overnight (37°C, 5% CO_2_). At minimum 4 hours before the experiment, the Seahorse XFe port injector sensor cartridge was placed in 200 µL of Seahorse XFe calibrant fluid. The day of the assay, culture media was aspirated and replaced with Seahorse XFe DMEM base media (1mM pyruvate, 2mM glutamine, 10mM glucose, pH 7.4) followed by incubation at 37°C, 0% CO_2_. Port injections were prepared in the same media and loaded into the cartridge with Port A containing test compound (serial diluted), port B containing oligomycin (1 µM post injection), port C containing FCCP (0.75 µM for MDA-MB-231, 0.58 µM for 4T1 post injection) and port D containing rotenone and antimycin A (0.5 µM each post injection). The port injection strategy was generated in Wave software, uploaded, the sensor cartridge and cell plate were inserted before the data collection.

### Seahorse XFe96 GlycoStress Test

MDA-MB-231 or 4T1 cells were seeded on a Seahorse XFe96 96-well plate at 20,000 cells/well and incubated overnight (37°C, 5% CO_2_). At minimum 4 hours before the experiment, the Seahorse XFe port injector sensor cartridge was placed in 200 µL of Seahorse XFe calibrant fluid. The day of the assay, culture media was aspirated and replaced with Seahorse XFe DMEM base media (2mM glutamine, pH 7.4) followed by incubation at 37°C, 0% CO_2_. Port injections were prepared in the same media and loaded into the cartridge with Port A containing test compound (serial diluted), port B containing glucose (10 mM post injection), port C containing Oligomycin (1 µM post injection) and port D containing 2-deoxyglucose (2-DG) (50mM post injection). The port injection strategy was generated in Wave software, uploaded, the sensor cartridge and cell plate were inserted before the data collection.

---

**Figure S1:**
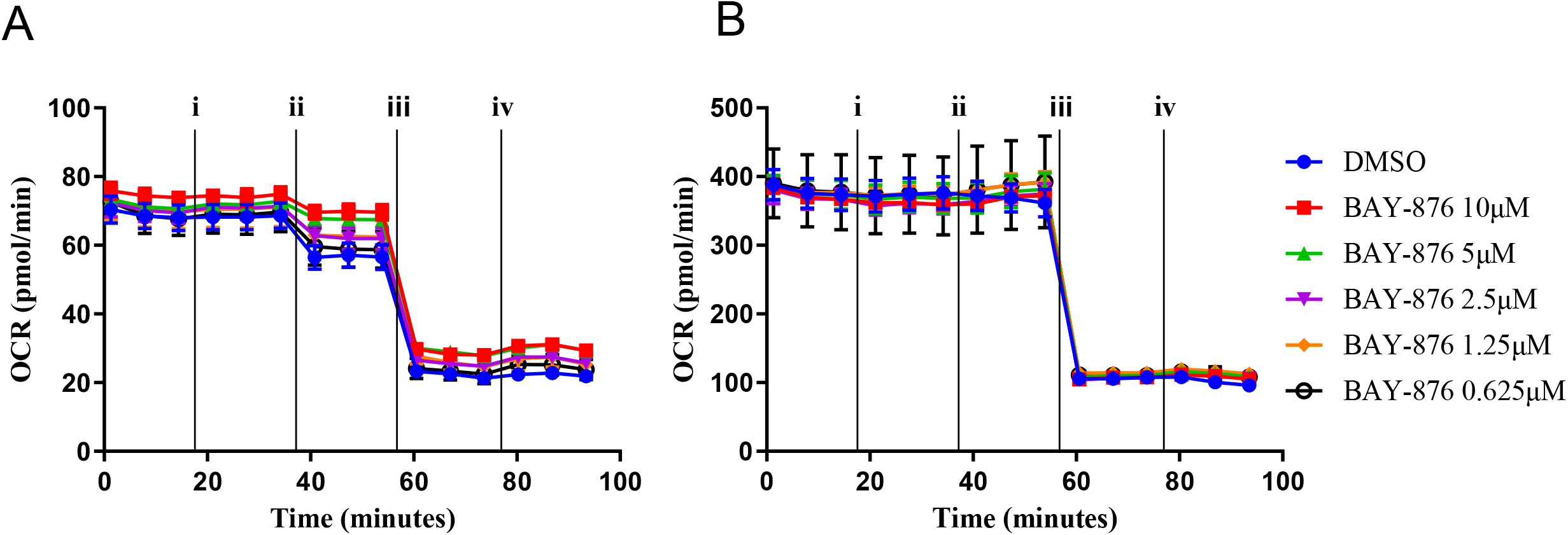
**A.** MDA-MB-231 oxygen consumption rates (OCR (pmol/min)) vs. time after injection of i) treatment or DMSO ii) glucose iii) oligomycin and iv) 2-deoxyglucose after treatment with BAY-876 (10-0.625µM). **B.** 4T1 oxygen consumption rates (OCR (pmol/min)) vs. time after injection of i) treatment or DMSO ii) glucose iii) oligomycin and iv) 2-deoxyglucose after treatment with BAY-876 (10-0.625µM).

**Figure S2:**
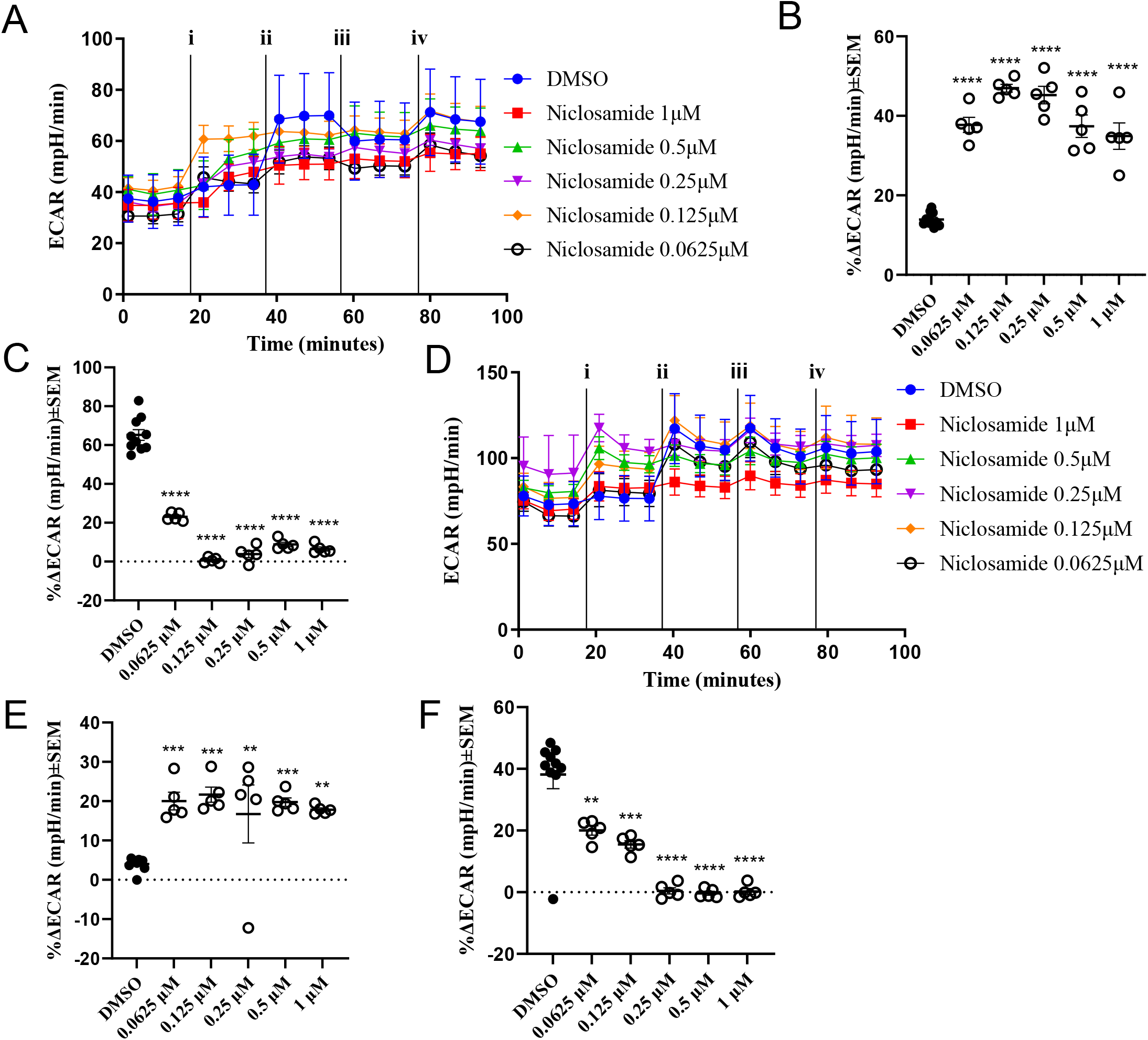
**A.** MDA-MB-231 extracellular acidification rates (ECAR (mpH/min)) vs. time after injection of i) treatment or DMSO ii) oligomycin iii) FCCP and iv) Rotenone and Antimycin A after treatment with niclosamide (1-0.0625µM). **B.** Percent change in ECAR after acute injection of niclosamide (1-0.0625µM) indicating compensatory glycolysis in MDA-MB-231 cells. **C.** Percent change in ECAR after injection of oligomycin indicating switch to glycolytic metabolism after uncoupling mitochondrial ATP production in MDA-MB-231 cells. **D.** 4T1 extracellular acidification rates (ECAR (mpH/min)) vs. time after injection of i) treatment or DMSO ii) oligomycin iii) FCCP and iv) Rotenone and Antimycin A after treatment with niclosamide (1-0.0625µM). **E.** Percent change in ECAR after acute injection of niclosamide (1-0.0625µM) indicating compensatory glycolysis in 4T1 cells. **F.** Percent change in ECAR after injection of oligomycin indicating switch to glycolytic metabolism after uncoupling mitochondrial ATP production in 4T1 cells. **B, C, E, F:** results are representative of 3 biological replicates experiments ± SEM. Statistical analysis was carried out via Dunnett’s one-way ANOVA compared to DMSO control (*p<0.05, **p<0.01, ***p<0.001, ****p<0.0001).

**Figure S3:**
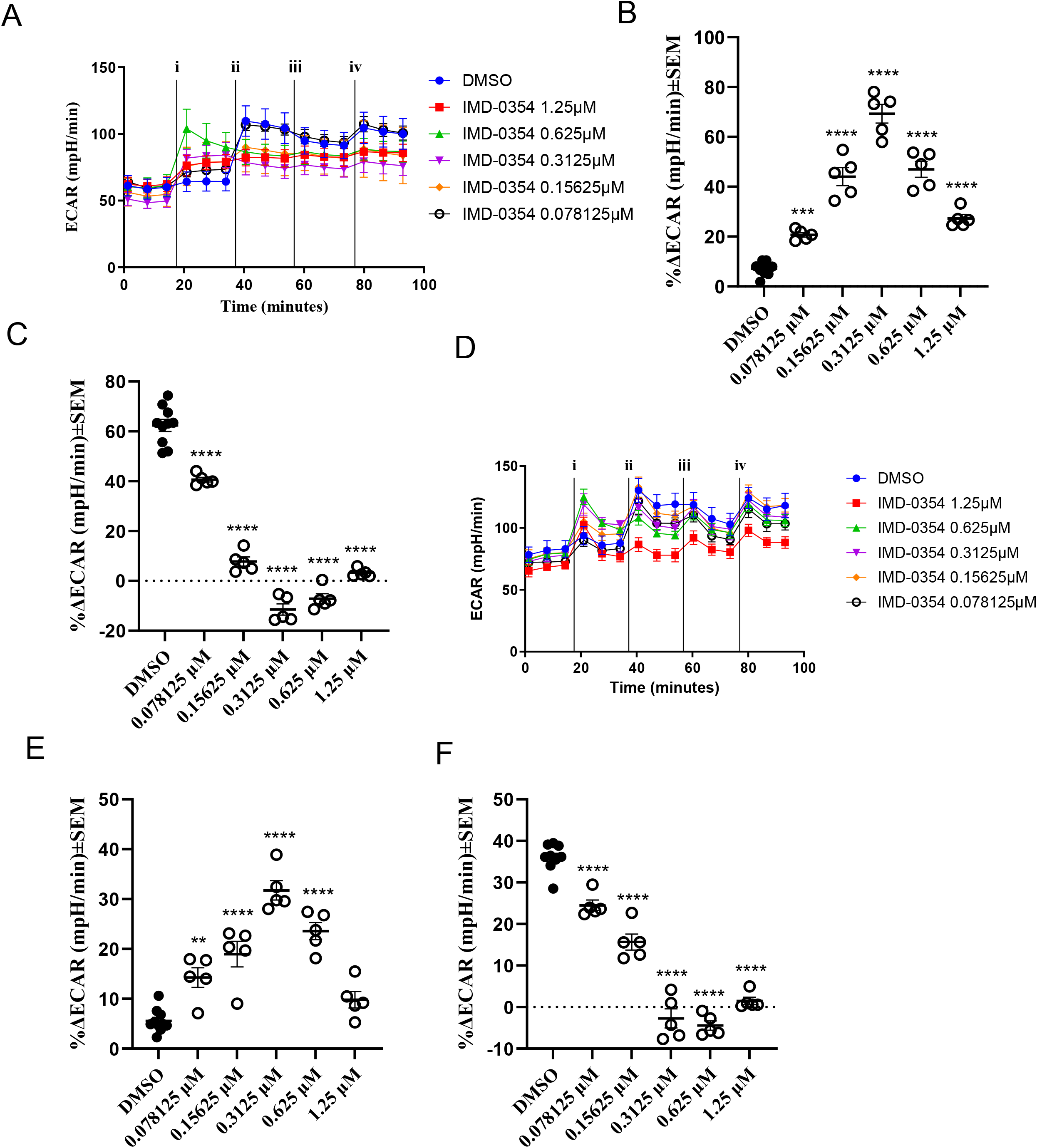
**A.** MDA-MB-231 extracellular acidification rates (ECAR (mpH/min)) vs. time after injection of i) treatment or DMSO ii) oligomycin iii) FCCP and iv) Rotenone and Antimycin A after treatment IMD-0354 (1.25-0.078125µM). **B.** Percent change in ECAR after acute injection of IMD-0354 (1.25-0.078125µM) indicating compensatory glycolysis in MDA-MB-231 cells. **C.** Percent change in ECAR after injection of oligomycin indicating switch to glycolytic metabolism after uncoupling mitochondrial ATP production in MDA-MB-231 cells. **D.** 4T1 extracellular acidification rates (ECAR (mpH/min)) vs. time after injection of i) treatment or DMSO ii) oligomycin iii) FCCP and iv) Rotenone and Antimycin A after treatment with IMD-0354 (1.25-0.078125µM). **E.** Percent change in ECAR after acute injection of IMD-0354 (1.25-0.078125µM) indicating compensatory glycolysis in 4T1 cells. **F.** Percent change in ECAR after injection of oligomycin indicating switch to glycolytic metabolism after uncoupling mitochondrial ATP production in 4T1 cells. **B, C, E, F:** results are representative of 3 biological replicates experiments ± SEM. Statistical analysis was carried out via Dunnett’s one-way ANOVA compared to DMSO control (*p<0.05, **p<0.01, ***p<0.001, ****p<0.0001).

**Figure S4:**
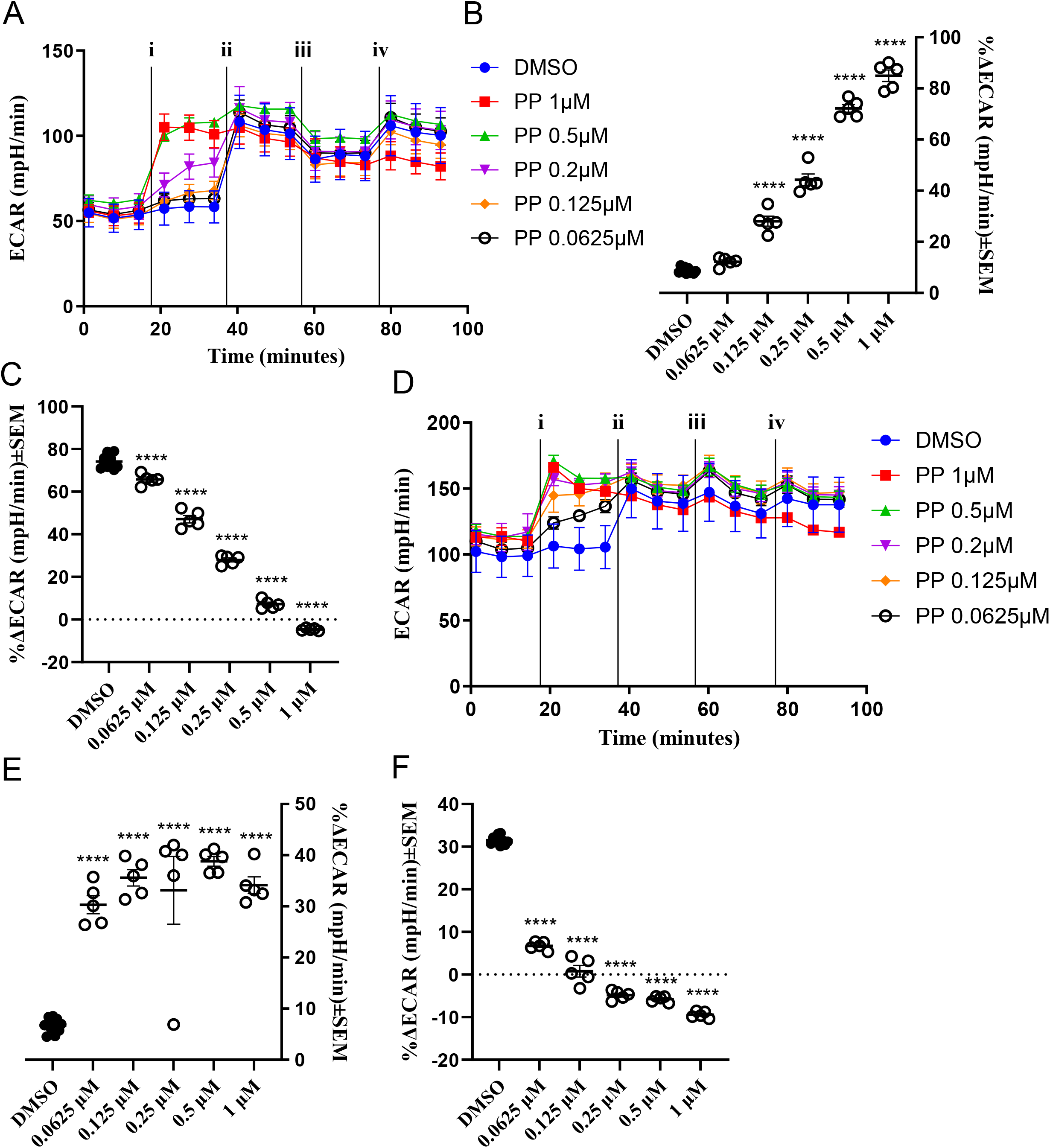
**A.** MDA-MB-231 extracellular acidification rates (ECAR (mpH/min)) vs. time after injection of i) treatment or DMSO ii) oligomycin iii) FCCP and iv) Rotenone and Antimycin A after treatment with PP (1-0.0625µM). **B.** Percent change in ECAR after acute injection of PP (1-0.0625µM) indicating compensatory glycolysis in MDA-MB-231 cells. **C.** percent change in ECAR after injection of oligomycin indicating switch to glycolytic metabolism after uncoupling mitochondrial ATP production in MDA-MB-231 cells. **D.** 4T1 extracellular acidification rates (ECAR (mpH/min)) vs. time after injection of i) treatment or DMSO ii) oligomycin iii) FCCP and iv) Rotenone and Antimycin A after treatment with PP (1-0.0625µM). **B.** Percent change in ECAR after acute injection of PP (1-0.0625µM) indicating compensatory glycolysis in 4T1 cells. **C.** percent change in ECAR after injection of oligomycin indicating switch to glycolytic metabolism after uncoupling mitochondrial ATP production in 4T1 cells. **B, C, E, F:** results are representative of 3 biological replicates experiments ± SEM. Statistical analysis was carried out via Dunnett’s one-way ANOVA compared to DMSO control (*p<0.05, **p<0.01, ***p<0.001, ****p<0.0001).

